# Thyroid Ablation Alters Passive Stiffness and Swimming Kinematics in Zebrafish

**DOI:** 10.1101/2022.09.09.507357

**Authors:** Pranav Parikh, Stacy Nguyen, Sarah McMenamin, Christopher P. Kenaley

## Abstract

Locomotion behavior is ultimately determined by the integration between active and passive tissues of an organism, but little is known about how these properties develop or are maintained. In this study, we used zebrafish (*Danio rerio*) to address the effects of a developmental hormone on morphogenesis and mechanical integration during swimming. We analyzed common kinematic variables and estimated intervertebral joint (IVJ) stiffness of zebrafish reared under different thyroid hormone profiles (euthyroid and hypothyroid) swimming during two different forward speeds, 5 and 10 BL·s^−1^. We found that zebrafish reared under hypothyroid conditions swam with higher trailing-edge amplitude, a larger amplitude envelope, longer propulsive wavelengths, and lower values of lateral strain in posterior regions at both speeds. IVJ second moment area about the bending axis was greater in the TH-, a result of a change in vertebral shape compared to wildtype fish. We conclude that thyroid hormone contributes to axial design during development and therefore has an important role in determining flexural stiffness and the swimming behaviors that are affected by this important property.

## Introduction

Most fishes swim by passing an undulating wave through the body that propagates rostrocaudally to produce thrust (Lighthill, 1971; Wardle et al., 1995; Blake, 2004). Understanding the kinematic and physiological contributions of undulatory thrust production amongst teleosts is a major focus research to this day (Shadwick & Gemballa, 2005; Lauder, 2015; Di Santo et al., 2021; Li et al., 2021). The shape and behavior of the undulatory wave in swimming fishes is the result of complex fluid-structure interactions (Hess & Videler, 1984; Tytell et al., 2010; Voesenek et al., 2020; Li et al., 2021). These interactions are, in turn, affected by the flexural stiffness along the body which itself is determined by the dynamics of muscle contraction and passive material properties of the body (Wakeling & Johnston, 1999; Tytell et al., 2010). While both active and passive properties of the musculoskeletal systems contribute to locomotor behavior, the processes that develop, regulate, and maintain these properties are poorly understood.

Circulating hormones, including androgens and estrogens, corticosteroids, insulin, and thyroid hormone, are crucial mediators of anatomical and physiological systems in vertebrates (Coughlin et al., 2001; Dugas et al., 2016; Keer et al., 2019; Pankova et al., 2019). Indeed, thyroid hormone (TH) stimulates major physiological, anatomical, and locomotor transformations during the metamorphic process in many vertebrates, including amphibians and teleosts (Kim & Mohan, 2013; McMenamin et al., 2017). In zebrafish and other developing teleosts, TH stimulates many components of metamorphosis from larva to juvenile (Power et al., 2001; McMenamin & Parichy, 2013; McMenamin et al., 2017; Hu et al., 2019). Importantly, TH is crucial in regulating skeletal development in fish, mediating skeletal transformations that create an overall adult body shape and physiology (Bassett & Williams, 2008; Shkil et al., 2014; Hur et al., 2017; Hu et al., 2019; Galindo et al., 2019; Keer et al., 2019, 2022). In rainbow trout, thyroid hormones induces a developmental shift to expression of faster functioning isoforms of the myosin heavy chain of myosin (Coughlin et al., 2001). Thus, TH mediates skeletogenesis and morphology, which contribute to passive flexibility, while also regulating metabolism and muscle performance, which likely contributes to passive and active properties of the body while swimming.

We hypothesize that developmental TH is required for typical integration of swimming kinematics. Specifically, we predict that the locomotor kinematic behavior will be altered in hypothyroid zebrafish (*Danio rerio*) as compared to wildtype counterparts. To evaluate this hypothesis, we pharmacologically ablated the thyroid system and studied the effect of this on key swimming-kinematic parameters including trailing-edge amplitude and frequency, propulsive wavelength, and amplitude envelope. Taken together, these parameters describe the speed and shape of the the propulsive wave. We also set out to evaluate whether axial muscle behavior and passive stiffness may differ between euthyroid and hypothyroid fish, differences that may contribute to altered kinematics. Therefore we assessed lateral strain, a proxy for axial muscle strain. To evaluate possible differences in passive stiffness, we calculated the second moment area and characterized the shape of intervertebral joints in specimens of both treatment groups.

## Methods

### Husbandry and Thyroid Ablations

For our hypothyroid group, we chose the transgenic Tg(tg:nVenus-v2a-nfnB) line (McMenamin et al., 2014). Transgenic thyroid ablations and control treatments were performed at 4 days post fertilization by placing larvae overnight in 10 mM metronidazole with 1% DMSO (to ablate the thyroid follicles for TH-), or with 1% DMSO alone (for euthyroid WT controls) in 10% Hanks, as in McMenamin et al. (2014). Fish were reared under standard conditions at 28°C with 14:10 light:dark cycle, and fed marine rotifers and Artemia pure Spirulina flakes (Pentair). For swimming trails, six age-matched and approximately size-matched (22–25 mm BL) males were taken from the TH- and WT groups and isolated from their respective populations. During the week of swimming trials, fish were kept in 23°C water and fed three times a week. For our analysis of second moment area of the intervertebral joints, we isolated an additional five individuals (23–25 mm mm BL) from the TH- and WT populations. Maintenance and experimentation of zebrafish fell under Boston College IACUC protocol 2017-007.

### Swimming Protocols and Kinematic Analysis

To avoid timing bias, trials alternated between TH- and WT individuals. All swimming experiments were conducted in a 28-L Brett-type tunnel (Loligo Systems) at room temperature (23±1°C). The working section of the swim tunnel measured (40 cm × 20 cm W × 20 cm D) and was modified by inserting a clear, 10-cm square acrylic tube to constrain the swimming position of the zebrafish specimens. Water velocity was controlled using a digital DC inverter (Eurodrive, Lyman) and calibrated inside the acrylic tube using a vane-wheel flow meter before each experiment. To ensure laminar, nonturbulent flow, 3D-printed ABS honeycomb was inserted in the upstream opening in the square tube. Each zebrafish was swam at two forward swimming speeds: 5 and 10 BL s^−1^. Trials alternated between speeds, with random assignment of which speed came first. Individual specimens were filmed from dorsal view during steady swimming over the course of 5 s with a Flea3 USB3 camera (FLIR Integrated Imaging Solutions) recording at 100 frames s^−1^ for a total of 500 frames. Each fish swam for between 3–5 seconds of the total recording, depending on individual and swimming flume speed. The procedure was repeated 3–5 times for a total recording time of 15–25 s at speed for each individual. Specimens were backlit with a 6-W LED light box placed below the clear bottom on the swim tunnel.

We analyzed frames representing a period in which the fish swam more than 0.5 cm away from the sides of the tube and was centered in the camera frame of view. To ensure that we analyzed only the steady-swimming phase of each sequence, we limited frames to those in which the fish changed its streamwise position by less than 2 mm (<0.08–0.9 BL) over the course of each tailbeat cycle. After applying these standards we amassed a total of 541 tailbeat cycles across the 12 fish for a mean of 45.1 (S.D.=18.0) for each.

To extract waveforms with high spatial and temporal resolution, we used the semiautomated tracking program trackter, written for the R computing environment (https://github.com/ckenaley/trackter). Briefly, tracker selects regions of interest by first converting images to binary format and then thresholding and segmentation. This method requires reasonable contrast between the ROI and background. Creating a silhouette with backlighting, a technique used in this study, is very effective in creating contrast (Beddow et al., 1995; Anderson & DeMont, 2000). Data returned from trackter include frame number, *x* and *y* positions of the head and tail, and amplitude of the tail. Midline data are also returned and include 200 *x* and *y* positions of the ROI midline and a smoothed *y* position of the midline. The midline of the body contour is calculated as the vertical midpoint between minimum and maximum *y*-pixel values at each *x*-pixel position along the body. Midline *x* and *y* data were passed through a LOEES smoothing operation to calculate a smoothed midline.

Using the smoothed midline data from the steady phase of swimming, we calculated maximum propulsive wavelength (*λ*, in BL) over each tailbeat cycle in the following manner. First, we used trackter to compute a theoretical horizontal midline established by a linear prediction of all points across the midline for each tailbeat cycle. This theoretical linear midline, which represents the axis of progression of the propulsive wave, was then subtracted from smoothed midline values to produce a standardized midline (*z*(*x*)) for each frame within a tailbeat cycle. Using this method, the longest full wavelengths over each tailbeat cycle were calculated as the distance between the internodes where the smooth midline intersects the axis of propulsive wave progression.

For each tail-beat cycle, we also calculated caudal-fin (i.e., trailing-edge) amplitude, frequency, and the amplitude envelope. Amplitude (in BL) and amplitude envelop (*z*(*x*) in BL) were computed as the maximum displacement of the last 2% of the smoothed midline and maximum lateral displacement along the body from the axis of progression, respectively. Frequency in Hz was computed as the inverse of each tail-beat cycle period.

To investigate TH’s role in muscle function, we also calculated maximum curvature of the body midline (*κ*(*x*) in BL) and, from this, peak lateral strain (*ϵ*(*x*), in BL) for each tail-beat cycle of steady swimming. Surface strain can be an accurate estimate of muscle strain (**?**Shadwick & Gemballa, 2005), thus we calculated *ϵ*(*x*) as the product of curvature*κ* and *h*(*x*), where *h*(*x*) is the body’s half thickness with respect to body position. Following Katz & Shadwick (1998), we calculated local curvature along the body as a function of the coordinate space defined by *x* and *z*, the longitudinal and lateral positions of the body midline in BL, respectively:

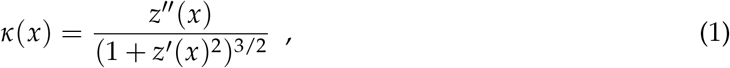

where *z*(*x*) is the function that describes the fish midline with respect to *x*, the longitudinal position, and is differentiated to obtain *z*′ and *z*″(*x*).

### Internal Passive Mechanics

The fish body may be treated as a simple beam, whereby the axial waveform is subject to the passive mechanics of the body (Long et al., 1994; Long & Nipper, 1996). Specifically, a wave passing through a body of relatively high flexural stiffness (*M*) will lengthen while one passing through low M will shorten. The bending stiffness of the axial skeleton as mediated by the properties of the intervertebral joints (IVJs) is particularly important in this regard. With their fluid-filled encapsulating membranes, IVJs may function as hydrostatic hinges that resist bending (Nowroozi et al., 2012; Nowroozi & Brainerd, 2012; Schmitz, 1995) To assess how passive axial mechanics may have been affected by our TH treatment and therefore affect wave propagation, we estimated the second moment area (*I*) of the 10 precaudal and 14 caudal intervertebral joints that span from just behind the pectoral girdle to the hypural region (25-80% BL) in five specimens from each treatment group.

Flexural stiffness is defined as *M* = *EI*, where *E* is the Young’s modulus. In this case, we must assume that *E* is relatively consistent throughout the intervertebral joints if *I* is to be a good approximation of their bending resistance. We note that the tissues within the intervertebral joints are heterogenous (Schmitz, 1995; Nowroozi et al., 2012; Nowroozi & Brainerd, 2012) and *E* likely varies accordingly; however, in their study of striped bass (*Morone saxatilis*,) Nowroozi et al. (2012) and Nowroozi & Brainerd (2012) showed that *I* of the intervertebral joints is a reasonably good predictor of joint flexibility.

We computed *I* of the intervertebral joints for each specimen from reconstructed transverse sections obtained through from *μ*-CT with a SkyScan1173 high-energy spiral-scan CT unit (MicroPhotonics, Inc.). Scan parameter values for amperage and voltage were 222 mA and 45 kV, respectively. Each scan produced a voxel size of 10 *μ*m. After slice reconstruction in NRecon (Micro Photonics, Inc.), we circumscribed the perimeter of the rostral centrum face of each of the precaudal and caudal vertebrae in imageJ (Schneider et al., 2012). These outlines, totaling 100 equally spaced *x-y* points each, were used to model the intervertebral joint of each vertebra as a solid ellipsoid whose second moment area about the y axis (*I_y_*) is described by

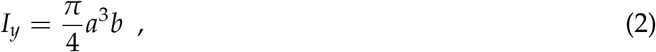

where *a* is the radius along the *x* axis and *b* is the radius along the y axis. The radii *a* and *b* were calculated as half the distance between the extrema of the *x* and *y* coordinates, respectively.

### Morphological Analysis

To visualize how vertebral shape contributes to *I*, we performed a principal components analysis (PCA) on the outlines of each joint, hypothesizing that wider IVJs will impart higher values of *I*. We first aligned the joint outlines with a full generalized procrustes alignment and then performed elliptic Fourier transformation (EFT) on the aligned outlines. EFT generates a number of shape-dependent variables, or harmonics, that are independent of size (Crampton, 1995). The number of harmonics was set to 8, a value that described 99% of the joint variation. Finally, we performed a PCA on the EFT coefficients and plotted the empirical IVJ morphospace based on the PC scores. Alignment, EFT, and PCA were all performed with the momocs package (Bonhomme et al., 2014) written for the R computing environment (R Core Team, 2022).

### Statistical Analysis

For trailing-edge amplitude and frequency, we assessed the difference between TH treatments using a mixed-effect linear model with specimen as a random effect and speed (5 or 10 BL s^−1^), thyroid treatment as a fixed effects and interactions for all terms. For amplitude envelope, propulsive wavelength, and lateral strain we employed the same mixed-effect linear models but also included body position as a covariate. During preliminary analysis, amplitude envelope appeared to increase exponentially with rostro-caudal body position, a near ubiquitous feature of swimming fishes (Di Santo et al., 2022). Therefore, in our linear mixed-effect model of amplitude envelope, the body position covariate was added as a second-order polynomial term. Lateral strain also appeared to follow a non-linear, but complex relationship with body position, and therefore the body position covariate was added as a polynomial spline term with the number of degrees of freedom, ranging from 3 to 10, chosen by Akaike Information Criterion weight (AICw). For our analysis of *I* at the IVJs, we also constructed a linear mixed-effect model, with specimen as a random effect, treatment as a fixed effect, and IVJ body position as a covariate.

The effect of speed, treatment, and any covariates in all models were evaluated with ANOVA. All statistical analyses were performed in R and considered significant at p<0.05.

## Results and Discussion

### Thyroid Status Regulates Important Kinematic Variables

Thyroid ablation and swimming speed had a significant effect on trailing-edge amplitude during the steading phase of swimming (both p<<0.001) and was markedly higher in TH- fish relative to WT controls across both speeds (Figure 2A). Thyroid condition had no effect on tail-beat frequency (p=0.311), nor did speed (p=0.189); however, we did uncover a significant interaction between speed and thyroid condition (p=0.001), indicating that the response of frequency to speed was different between the two treatment groups. Specifically, frequency was higher at 10 BL·s^−1^ than at 5 BL·s^−1^ in our wild-type, euthyroid group, but frequency remain constant between the two speeds in the ablated group (Figure 2B). Speed and treatment were significant predictors of maximum propulsive wavelength (both p¡0.001) with higher *λ* at 10 BL·s^−1^ than at 5BL·s^−1^ and in the ablated group (Figure 2C).

We found the expected pattern of increasing amplitude envelope with respect to body position described by a second order polynomial term (p<<0.001). We identified a significant interaction between treatment, speed, and body position (p<<0.001), with the envelope greater at 10 BL·s^−1^ than at 5 BL·s^−1^ in both treatment groups and *z*(*x*) increasing more along the body in TH- fish from 5 BL·s^−1^ to 10 BL·s^−1^ compared to WT (Figure 1A).

**Figure 1:**
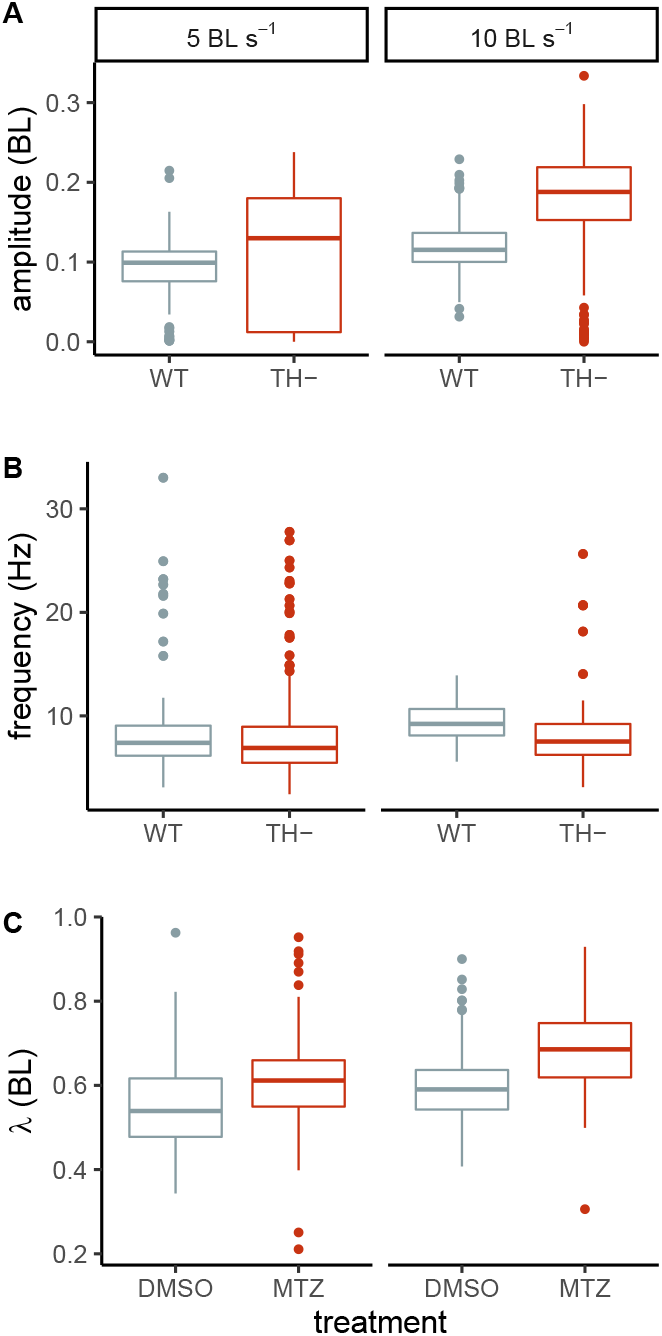
Boxplots comparing (A) trailing edge frequency, (B) amplitude, and (C) maximum propulsive wavelength of wild-type (“WT”) and thyroid-ablated (“TH-”) zebrafish swimming at two speeds, 5 and to 10 BL·s^−1^.

For maximum lateral strain, we found a polynomial spline term for body position with 10 degrees of freedom fit best (AICw= 1.00) and that this model predicts a significant increase along the body (p<<0.001), peaking at about 0.8 BL and decreasing sharply thereafter in both treatment groups and speeds (Figure 1B). We also found a significant interaction between treatment, speed, and body position (p<<0.001). There was a marked contrast in the relative increase of lateral strain with body position between the two treatment groups in that the thyroid-ablated group underwent a lesser degree of strain in the posterior of the body at both speeds and the magnitude of change in maximum curvature in the posterior from 5 BL·s^−1^ to 10 BL·s^−1^ was greater in the WT group (Figure 1B).

In all, our kinematic results show that WT and TH- individuals generally followed canonical patterns of fish swimming kinematics. Both groups exhibit typical patterns of an increasing amplitude envelope and magnitudes of lateral strain and lateral displacement along the body and increased magnitudes of strain at faster speeds (Di Santo et al., 2021).

There were, however, conspicuous differences in several kinematic parameters between the thyroid treatment groups; some of these difference suggest abnormal swimming mechanics in TH- fish. Fishes typically attain faster forward swimming speeds by modulating tail-beat frequency (Lighthill, 1971; Li et al., 2021). The ablated group showed no increase in frequency at a faster speed, while the tail-beat frequency of the euthyroid group follows the typical pattern (Figure 2B). Consequently, tail-beat frequency was lower in TH- fish at 10 BL·s^−1^. Tail-beat amplitude in the TH- group was higher than in the eurthyroid group at both speeds and highest at 10 BL·s^−1^. Maximum propulsive wavelength increased from 5 BL·s^−1^ to 10 BL·s^−1^ while it stayed relatively constant at the higher speed int WT fish. Thus, it appears speed modulation in the TH- group was mediated by changes of greater magnitude in amplitude rather than frequency and that this change is related to mechanics associated with changes in propulsive wavelength. Furthermore, the magnitude of lateral strain in posterior regions of the body were much lower in the TH- group than in WT fish.

**Figure 2:**
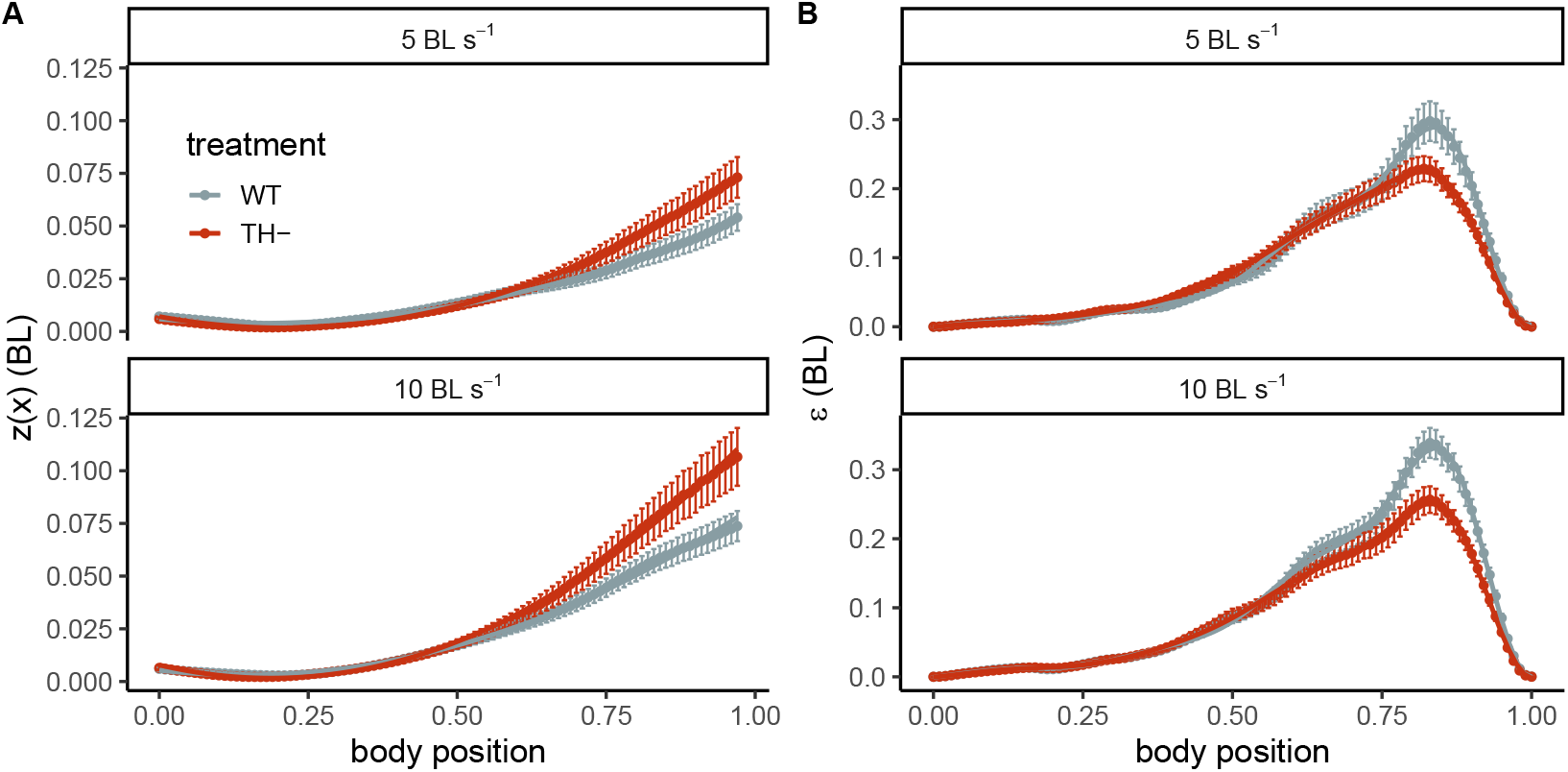
The (A) amplitude envelope (z(x) and (B) maximum lateral strain (*ϵ*) along the body of wild-type (“WT”) and thyroid-ablated (“TH-”) zebrafish swimming at two speeds, 5 and to 10 ·s^−1^. Mean values are represented at each 0.01 BL position ± 10 S.E.M. Trends according to treatment and speed in A were fit with a second order polynomial term for the body position covariate, while trends in B were fit with a 10^*th*^ order polynomial spline term.

### Thyroid Status Alters Vertebral Morphology and Passive Stiffness

Excess thyroid hormone is known to specifically affect vertebral morphology in other cypriniform species (Bolotovskiy & Levin, 2015) and causes vertebral demineralization in eels (Sbaihi et al., 2007). We therefore tested whether thyroid status affected the morphology of IVJs in zebrafish, and whether this could contribute to a change in passive body stiffness.

The first two PCs of our PCA analysis based on EFT captured 84.3% of IVJs shape variance (PC=78.4%, PC2=5.87%). PC1 represented a change in aspect ratio, with higher PC scores representing wider IVJs and lower scores representing more round IVJs. TH- fish occupied areas of wide morphospace, while WT fish occupied areas of round morphospace (Figure 3A). The areas of morphospace representing the widest IVJs were occupied by caudal vertebrae of the ablated fish. Thyroid status also showed a significant interaction with body position in predicting *I* of the IVJs (p<<0.001): while *I* was relatively constant along the body in wild-type fish, *I* increased with body position in the TH- group (Figure 3B). Mean second moment area for TH- fish was 1.5 times greater in the precaudal IVJs and 3.4 times great in the caudal IVJs compared to WT fish.

**Figure 3:**
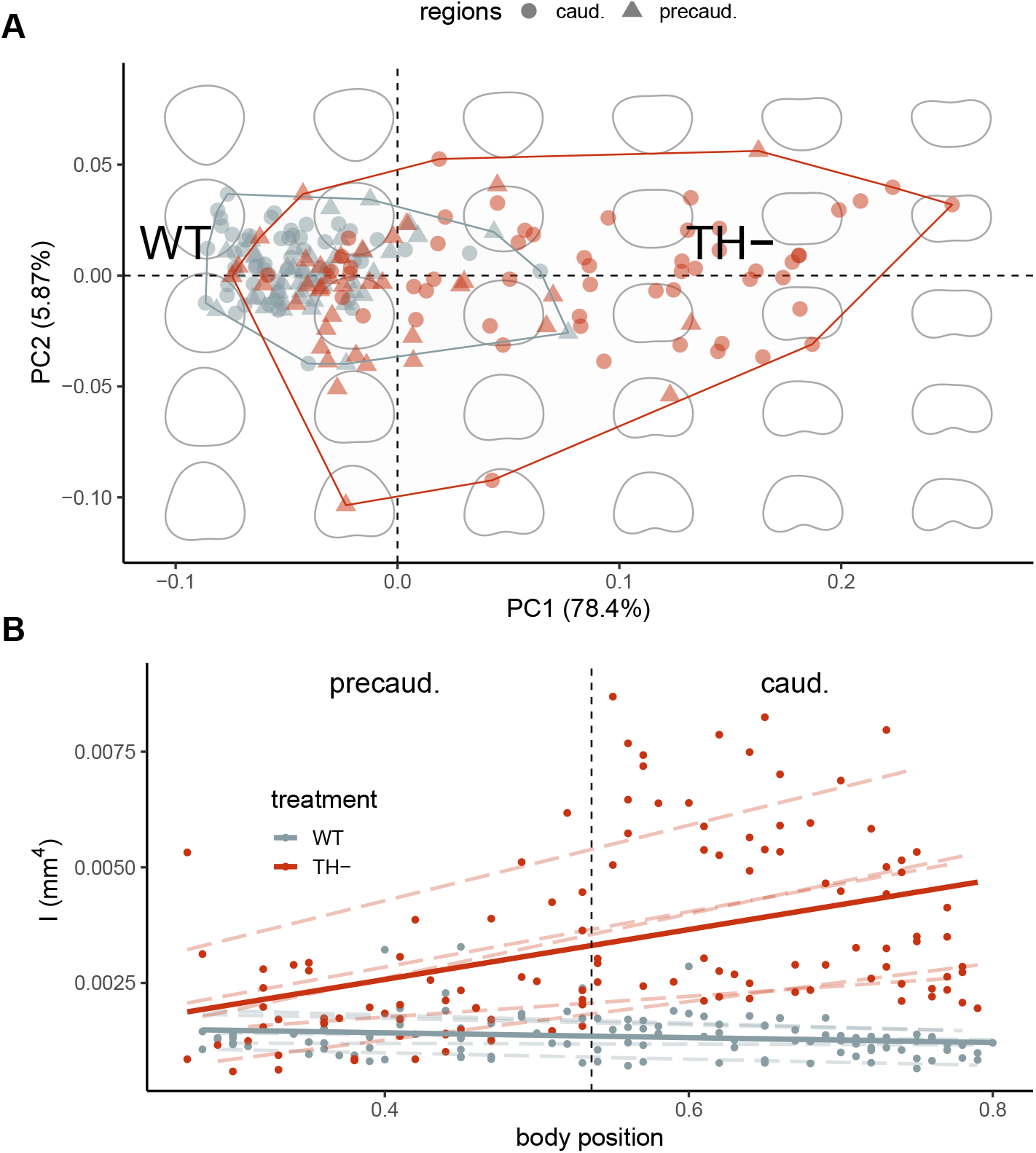
(A) a PCA biplot and morphospace of intervertebral joints of wild-type (“WT”) and thyroid-ablated (“TH-”) zebrafish and (B) the relationship of second moment area (*I*) of intervertebral joins (IVJs) along the body. Symbols in A correspond to either precaudal or caudal IVJs. Outlines in the background of A represent morhpospace positions described by the first two principal components of an analysis based on elliptical Fourier transformation. Dashed trend lines in B represent linear trends for each specimen (n=5 for each treatment), while the solid lines represent linear trends for each treatment. The dashed vertical line in B represents the division along the body length between predcaudal and caudal IVJs.

In summary, we have found that higher *I* with reference to the bending axis about *y* in the treatment group, particularly in caudal IVJs, is the logical outcome of a redistribution of IVJ area along the mediolateral (*x*) axis of the transverse plane. By altering IVJ morphology in such a way, vertebrae of TH- fish show an axial skeleton with functionally greater passive stiffness.

### Altered Kinematics with Passive Stiffness

To our knowledge, this study is the first to developmentally alter the passive stiffness of a fish and investigate the resultant changes in locomotor behavior. Our results underscore the important and complex interactions between passive mechanics and swimming kinematics in the production of locomotor thrust. Undulatory swimmers produce thrust to overcome hydrodynamic drag during propulsion; long-standing locomotor theory posits that fishes produce the requisite amount of thrust at a given speed by through a combination of entraining enough water over a tailbeat cycle (i.e., a large lateral displacement) and entraining water at the appropriate rate (i.e., a certain driving frequency; Bainbridge, 1958; Lighthill, 1971; Sfakiotakis et al., 1999; Di Santo et al., 2021). By extension, fishes may entrain more water through increased lateral displacement or increased driving frequency to increase locomotor speed. Our results demonstrate that the tradeoff between increasing amplitude or frequency in the modulation of speed can be altered by the passive stiffness of the body. Specifically, (1) a stiffer body relies on longer wavelengths of higher amplitudes to produce the requisite amount of thrust at a given speed, whereas (2) a less stiff body relies on higher higher driving frequencies of lower amplitudes.

Other authors have, through indirect means, demonstrated the importance of passive stiffness in mediating swimming performance. Experimental manipulations of body stiffness and the study of physical models have uncovered the role passive stiffness has in achieving higher swimming speeds through increased tailbeat frequency, tailbeat amplitude, or propulsive wave-length (McHenry et al., 1995; Long et al., 1996, 2004, 2006; Esposito et al., 2012). Here, we have used thyroid ablation to directly alter the stiffness of a fish body, showing that a stiffer body results in higher trailing-edge amplitude, a larger amplitude envelope, and longer propulsive wavelengths at at higher speeds, directly corroborating the predictions of these previous studies.

We note again that our conclusions are predicated on *I* of the IVJs representing flexural stiffness (*EI*) of the axial skeleton. For this, we must also assume that *E* is invariant throughout the IVJs. The tissues within the intervertebral joints are indeed heterogenous (Schmitz, 1995; Nowroozi et al., 2012) and thus *E* likely varies. However, in their study of *Morone saxatilis*, Nowroozi et al. (2012) and Nowroozi & Brainerd (2012) revealed that *I* is good predictor of IVJ flexural stiffness.

Lastly, active, as well as passive components of the fish body contribute to overall body stiffness. The amount to which fishes actively modulate body stiffness with their axial musculature has been a fundamental thrust of research in undulatory propulsion (McHenry et al., 1995; Nowroozi & Brainerd, 2014; Jusufi et al., 2017). Dynamic tuning of body stiffness has been hypothesized to substantially reduce the internal resistance to bending by matching the body’s natural and driving frequencies, which in turn reduces the force required to bend the body at a given frequency (Long, 1998). In addition, contralateral muscle contractions could result in higher speeds as they store and return more energy to the water (McHenry et al., 1995). Our experimental approach and results cannot directly address how muscle activation pattern and changes in active stiffness are affected by thyroid status or whether such changes affect swimming behavior.

The absence of thyroid hormone during development may certainly affect muscle dynamics during swimming in zebrafish. In rainbow trout, flounder and rats, thyroid hormones induce a developmental shift in expression of different isoforms of important fiber proteins, including, troponin or myosin light and heavy chains of myosin (Gambke et al., 1983; Butler-Browne et al., 1984; Yamano et al., 1991; Coughlin et al., 2001). In trout, Coughlin et al. (2001) found that thyroid treatments also alter the contraction time and duty cycle of axial muscles. If thyroid-ablated TH- zebrafish possess inappropriate muscle activation patterns or lack a shift to developmentally appropriate isoforms of contractile elements, their ability to modulate speed through increased driving frequencies could be compromised, resulting from a reliance on poorly tuned muscles to bend a stiffer body. Our lateral strain results could potentially be attributed to aberrant tuning of force-production at high tail-beat frequencies.

## Acknowledgments

We wish to thank the members of the Kenaley and McMenamin Labs, past and present, as well as the staff of the BC Animal Care Facility.

## Author Contributions

PP and CK performed the experiments, captured kinematic data, and performed the analysis. CK and SM conceived the study. CK, PP, and SM helped write the manuscript and SN undertook *μ*-CT analysis.

## Supplementary Information

### Data Availability

Kinematic and morphological data and the R script used in analysis are archived on Dryad (`doi:10.5061/dryad.6m905qg34`). The R package trackter used in kinematic analysis is available at https://github.com/ckenaley/trackter.

